# R code and downstream analysis objects for the scRNA-seq atlas of human breast spanning normal, preneoplastic and tumorigenic states

**DOI:** 10.1101/2021.11.30.470523

**Authors:** Yunshun Chen, Bhupinder Pal, Geoffrey J Lindeman, Jane E Visvader, Gordon K Smyth

**Author notes:** corresponding author: Gordon Smyth.

## Abstract

Breast cancer is a common and highly heterogeneous disease. Understanding the cellular diversity in the mammary gland and its surrounding micro-environment across different states can provide insight into the cancer development in human breast. Recently, a large-scale single-cell RNA expression atlas was constructed of the human breast spanning normal, preneoplastic and tumorigenic states. Single-cell expression profiles of nearly 430,000 cells were obtained from 69 distinct surgical tissue specimens from 55 patients. This article extends the study by providing downstream processed R data objects, complete cell annotation and R code to reproduce all the analyses. Details of all the bioinformatic analyses that produced the results described in the study are provided.

## Background & Summary

Breast cancer is the most commonly diagnosed cancer and the leading cause of cancer death in women^1^. It is a very heterogeneous disease at the molecular level^2^. Different breast cancer subtypes can be characterized on the basis of expression profiles of markers such as estrogen receptor (ER), progesterone receptor (PgR), and human epidermal growth factor receptor 2 (HER2)^3^. The development of certain cancer subclasses is also known to be associated with mutations such as BRCA1^4^. Recently, we and colleagues constructed a large-scale single-cell RNA expression atlas of the human breast spanning normal, preneoplastic and tumorigenic states (subsequently referred to as the ScBrAtlas)^5^. Single-cell expression profiles of nearly 430,000 cells were obtained from 69 distinct surgical tissue specimens from 55 patients (Figure 1). This article extends the ScBrAtlas by providing downstream processed R data objects, complete cell annotation and R code to reproduce all the analyses.

**Figure 1.**
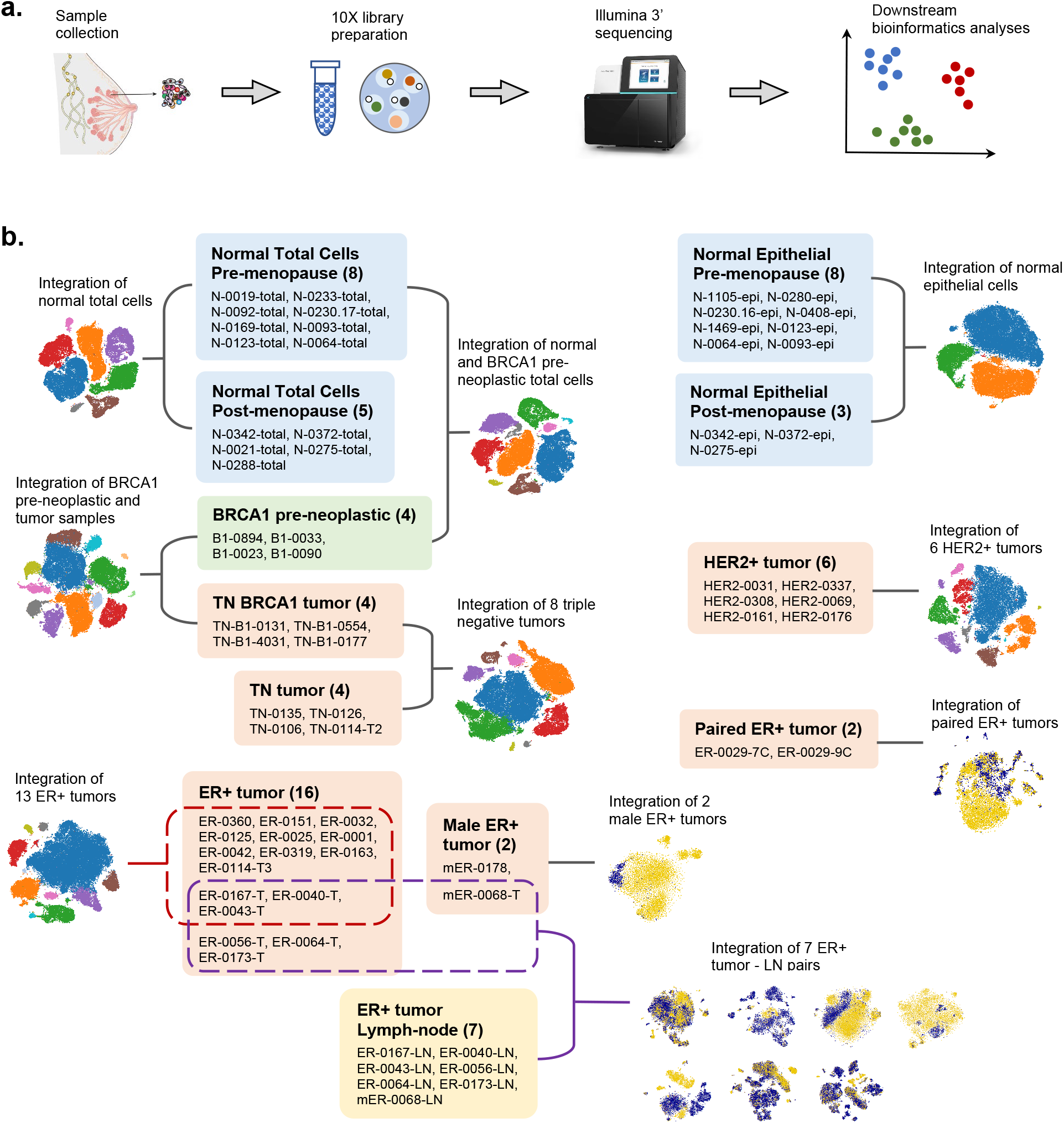
(a) Diagram showing the data processing pipeline from sample collection to downstream bioinformatics analyses. (b) Schematic overview of the all the integration analyses and the samples involved in each integration analysis. Under each category, the names of the samples are listed and the total number of samples is shown in the bracket.

The ScBrAtlas spanned several stages of breast cancer genesis. First, reduction mammoplasties were obtained from women with no family history of breast cancer to explore cellular diversity in normal breast epithelia as well as complexity within the normal breast ductal micro-environment. Three major epithelial cell populations revealed in literature^6^: basal, luminal progenitor (LP), and mature luminal (ML), were confirmed by the bulk RNA-seq signatures for sorted epithelial populations as well as the cell clustering of the integrated single cell transcriptomic data on normal breast epithelia. Similar cell type composition within the normal epithelium was observed across multiple healthy donors with different hormonal status (pre- and post-menopausal). For the immune and stromal micro-environment of normal breast tissue, integration analysis and the pseudo-bulk differential expression analysis identified different cell clusters including fibroblasts, endothelial cells (vascular and lymphatic), pericytes, myeloid, and lymphoid cells. Differential abundance analysis revealed that fibroblasts are more abundant whereas vascular endothelial cells are less abundant in post-menopausal tissue compared to pre-menopausal tissue^5^.

Next, breast tissue from BRCA1 mutation carriers was obtained for investigating cellular changes in precancerous state. Overall, the differences of stromal and immune subsets between normal and BRCA1+/− preneoplastic tissue were not significant, nor was the proportions of different cell clusters. However, extensive changes in the tissue micro-environment were observed between the preneoplastic and the neoplastic states in BRCA1 mutation carriers^5^.

Finally, ER+, HER2+ and triple negative breast cancer (TNBC) tumors were obtained from treatment-naive patients for exploring the degree of heterogeneity within the cancer cell compartment and its micro-environment across different tumor subtypes. Extensive inter-patient heterogeneity was revealed by single cell integration analyses across all cancer subtypes. Within the tumor populations, a discrete cluster of cycling MKI67+ tumor cells were observed for all three major breast cancer subtypes. Within the tumor micro-environment, different immune landscapes were observed in different cancer subtypes. Both TNBC and HER2 featured a proliferative CD8+ T-cell cluster, whereas ER+ tumors primarily comprised cycling TAMs. In addition, matched pairs of ER+ tumors and involved lymph nodes were profiled for examining the relationship between primary breast tumors and malignant cells that seed lymph nodes. Clonal selection and expansion were observed in some patients, whereas mass migration of cells from the primary tumor to the LN was observed in some other patients^5^.

The ScBrAtlas provides a valuable resource for understanding cellular diversity and cancer genesis in human breast. The examination and exploration of the single cell data presented in this study required large-scale bioinformatics analyses for multiple groupings of the original data. While genewise read counts were previously made publicly available for all 421,761 individual cells^7^, downstream results after quality filtering, data integration and cell clustering were not provided.

In this report we describe the bioinformatics analysis used in the ScBrAtlas in greater detail. We provide a complete description of the quality control filters used to select 341,874 cells for downstream analyses. The technical quality of both the 10X single-cell transcriptomic data sets and the bulk RNA-seq reference data set is assessed to demonstrate the reliability of the data. We provide downstream R data objects corresponding to each data integration and cell clustering presented in the ScBrAtlas, together with R code to reproduce the data objects. Crucially, the data objects provided here include cell barcodes by which each individual cell can be tracked through all the analyses. We also provide detailed information allowing the copy number variation analyses to be mapped back to individual samples and cell clustered, thus providing a way to distinguish putative malignant cancer cells from normal epithelial cells in the cancer tumors. All the resources and the detailed information can be easily accessed and utilized by researchers for further exploration and clinical validation, which may lead to discoveries of novel approaches for personalized breast cancer treatment in the future.

## Methods

### Read alignment and count quantification

Single-cell RNA-seq expression profiles of 69 samples from 55 patients were generated by the 10x Genomics Chromium platform and an Illumina NextSeq 500 system (Fig. 1a, Supplementary Table 1). The original Illumina BCL output was converted to FASTQ files and then aligned to the human reference genome GRCh38 (cellranger ref v3.0.0) using Cell Ranger v3.0.2 (https://support.10xgenomics.com). The outputs for each individual sample contain a count matrix in matrix market mtx.gz format, barcode information and feature information both in tab-delimited tsv.gz format (Supplementary Table 1). Any cell with at least 500 sequence reads assigned to genes was included in this output. All the downstream bioinformatics analyses were performed based on the cellranger outputs.

### Quality control and cell filtering

Sequence read counts were obtained for a total of 421,761 cells across the 69 samples (Supplementary Table 2). Quality control (QC) was performed individually for each scRNA-seq sample. Cells with high proportion of mitochondrial reads were considered as of low quality and hence were filtered^8^. A lower bound of 500 was generally applied to the number of detected genes for each cell, although this was reduced to 400 or 300 for a small number of samples with low read coverage. Upper bounds of a combination of number of detected genes and library size were also applied to each sample to remove potential doublets. The threshold values of these QC metrics for each individual sample are shown in Supplementary Table 2 and are also supplied in machine-readable form as part of this data submission^9^. A total of 341,874 cells remained after quality filtering for downstream analysis.

### Single-cell RNA-seq integration analysis

The samples included breast tissues from normal healthy donors, BRCA1 mutation carriers and patients diagnosed with different types of breast cancer (triple negative, ER+ and HER2+). Matching pairs of tumor and lymph node (LN) samples, as well as tumor samples from male patients, were also included. The single-cell analysis strategy involved grouping together comparable samples, integrating the profiles, then clustering cells into putative cell types. A total of 16 different sample-groups were integrated (fig. 1b). Some samples were involved in more than one integration, for example the pre-neoplastic samples with BRCA1 mutations were integrated first with the normal samples and later with the BRCA1 triple negative (TN) tumor samples. For some sample-groups analyses, subsets of cells were extracted, re-integrated and re-clustered. The total number of cell cluster analyses is shown in Table 1.

**Table 1.** Cell cluster analyses. Each row corresponds to one integration and cell clustering, except for TumLN, where one clustering was done for each of the 7 patients. Columns indicate the group of samples integrated, the cell subset clustered and the figure reference in the original ScBrAtlas study^5^.

| Label | Tissue Sample Type | Cell Family | Figure |
| --- | --- | --- | --- |
| NormEpi | Normal breast | epithelial cells | EV1C |
| NormEpiSub | Normal breast | epithelial cells without stroma | 1E |
| NormTotal | Normal breast | total cells | 2B |
| NormTotalSub | Normal breast | non-epithelial | 2D |
| NormTotalFib | Normal breast | fibroblast cells | 3D |
| NormB1Total | Normal and BRCA1 preneoplastic | total cells | 4B |
| NormB1TotalSub | Normal and BRCA1 preneoplastic | non-epithelial | 4C |
| BRCA1Tum | BRCA1 preneoplastic and BRCA1 TNBC | total cells | 4E |
| BRCA1TumSub | BRCA1 preneoplastic and BRCA1 TNBC | non-epithelial | 5A |
| TNBC | TNBC | total cells | 6A |
| HER2 | HER2+ breast tumor | total cells | 6B |
| ERTotal | ER+ breast tumor | total cells | 6C |
| ERTotalTum | ER+ breast tumor | epithelial cells | 6E |
| PairedER | Two ER+ breast tumors from patient 0029 | total cells | 6H |
| TNBCSub | TNBC | non-epithelial | 7A |
| HER2Sub | HER2+ breast tumor | non-epithelial | 7B |
| ERTotalSub | ER+ breast tumor | non-epithelial | 7C |
| TNBCTum | TNBC | epithelial cells | EV3B (top) |
| HER2Tum | HER2+ breast tumor | epithelial cells | EV3B (bottom) |
| TNBCTC | TNBC | T-cells | EV4A (left) |
| HER2TC | HER2+ breast tumor | T-cells | EV4A (middle) |
| ERTotalTC | ER+ breast tumor | T-cells | EV4A (right) |
| Male | ER+ breast tumors from male patients | total cells | EV5A |
| TumLN | ER+ breast tumor & lymph-node pairs from 7 patients | total cells | 9A |

Samples were integrated using the Seurat anchor-based integration method^10^. To perform dimensionality reduction, the first 30 principal component were computed and used for the cell clustering and t-distributed stochastic neighbor embedding (t-SNE) visualization^11^. The default Louvain clustering algorithm^12^ was used for cell cluster identification. Different resolution parameters were used in different cell clustering analyses to ensure repeatability and the best interpretation of the data^5^.

We provide here the Seurat data objects containing each of the cluster analyses as R data files (Table 2). The R data objects contain cell cluster details for each cell. The R code by which each R object was constructed is also provided (Table 2).

**Table 2.** Files deposited on Figshare^9^. Data files are in RDS format. Each data file contains one Seurat object except for TumLN, which contains a list of 7 Seurat objects. Each Seurat data object provides cell cluster identities and associated information for the corresponding cell cluster analysis. Code files contain the R code used to produce the corresponding Seurat objects.

| Label | Data filename | Code filename |
| --- | --- | --- |
| NormEpi | SeuratObject_NormEpi.rds | NormEpi.R |
| NormEpiSub | SeuratObject_NormEpiSub.rds | NormEpi.R |
| NormTotal | SeuratObject_NormTotal.rds | NormTotal.R |
| NormTotalSub | SeuratObject_NormTotalSub.rds | NormTotal.R |
| NormTotalFib | SeuratObject_NormTotalFib.rds | NormTotal.R |
| NormB1Total | SeuratObject_NormB1Total.rds | NormBRCA1.R |
| NormB1TotalSub | SeuratObject_NormB1TotalSub.rds | NormBRCA1.R |
| BRCA1Tum | SeuratObject_BRCA1Tum.rds | BRCA1Tum.R |
| BRCA1TumSub | SeuratObject_BRCA1TumSub.rds | BRCA1Tum.R |
| TNBC | SeuratObject_TNBC.rds | TNBC.R |
| TNBCSub | SeuratObject_TNBCSub.rds | TNBC.R |
| TNBCTC | SeuratObject_TNBCTC.rds | TNBC.R |
| TNBCTum | SeuratObject_TNBCTum.rds | TNBC.R |
| HER2 | SeuratObject_HER2.rds | HER2.R |
| HER2Sub | SeuratObject_HER2Sub.rds | HER2.R |
| HER2TC | SeuratObject_HER2TC.rds | HER2.R |
| HER2Tum | SeuratObject_HER2Tum.rds | HER2.R |
| ERTotal | SeuratObject_ERTotal.rds | ER.R |
| ERTotalSub | SeuratObject_ERTotalSub.rds | ER.R |
| ERTotalTC | SeuratObject_ERTotalTC.rds | ER.R |
| ERTotalTum | SeuratObject_ERTotalTum.rds | ER.R |
| Male | SeuratObject_Male.rds | Male.R |
| PairedER | SeuratObject_PairedER.rds | PairedER.R |
| TumLN | SeuratObject_TumLN.rds | TumLN.R |

### Differential expression and pathway analysis

Differential expression analyses were performed to detect marker genes for different cell clusters. In order to account for the biological variation between different patients, a pseudo-bulk approach was used in most cases where read counts from all cells under the same cluster-sample combination were summed together to form pseudo-bulk samples. The edgeR’s quasi-likelihood pipeline was used for pseudo-bulk differential expression analysis, where the baseline differences between patients were incorporated into the linear model^13^. The Seurat’s FindMarkers function was applied where pseudo-bulk samples were not satisfactory due to low cell numbers or imbalanced cluster-sample combination. KEGG pathway analyses were performed using the kegga function of the limma package^14^.

### Data visualization

Ternary plot visualization was performed as previously described^15^. Ternary plots position cells according to the proportion of basal, LP- or ML-positive signature genes expressed by that cell and were generated using the vcd package^16^. The t-SNE visualization for all the integration analyses were generated using the RunTSNE function in Seurat with a random seed of 2018 for reproducibility. Diffusion plots were generated using the destiny package^17^. MDS plots were created with edgeR’s plotMDS function. Log2-CPM values for each gene across cells were calculated using edgeR’s cpm function with a prior count of 1. Heat maps were generated using the pheatmap package. Log2-CPM values were standardized to have mean 0 and standard deviation 1 for each gene before producing the heat maps, after which genes and cells were clustered by the Ward’s minimum variance method^18^.

### Bulk RNA-seq data and differential expression analysis

RNA-seq experiments were performed to obtain signature genes of basal, luminal progenitor (LP), mature luminal (ML) and stromal cell populations. Epithelial cells for basal, LP, and ML populations were sorted from eight independent patients and stroma from five patients. For one particular patient, samples were collect from both left and right breast for each of the four cell populations. For another patient, ML cell population was collected twice. The complete RNA-seq data contains 9 basal, 9 LP, 10 ML and 6 stroma samples. RNA-seq libraries were prepared using Illumina’s TruSeq protocol and were sequenced on an Illumina NextSeq 500.

Reads were aligned to the hg38 genome using Rsubread v1.5.3^19^. Gene counts were quantified by Entrez Gene IDs using featureCounts and Rsubread’s built-in annotation^20^. Gene symbols were provided by NCBI gene annotation dated 29 September 2017. Immunoglobulin genes as well as obsolete Entrez Ids were discarded. Genes with count-per-million above 0.3 in at least 3 samples were kept in the analysis. TMM normalization was performed to account for the compositional biases between samples.

Differential expression analysis was performed using limma-voom^21^. Patients were treated as random effects and the intra-patient correlation was estimated by the duplicateCorrelation function in limma. Pairwise comparisons between the four cell populations were performed using TREAT with a fold change threshold of 1.5^22^. An FDR cut-off of 0.05 was applied for each comparison. Genes were considered as signature genes for a particular cell type if they were upregulated in that cell type in all the pairwise comparisons. The analysis yielded 515, 323, 765, and 1094 signature genes for basal, LP, ML, and stroma, respectively. In this submission we provide gene symbols of the signature genes as an R data file and R code to reproduce the bulk RNA-seq analysis^9,23^

### Differential abundance analysis

Differential abundance analyses were performed to examine the differences in cell cluster frequencies between pre-menopause and post-menopause groups in normal breast tissue micro-environment. Quasi-multinomial and quasi-binomial generalized linear models were used in order to account for the inter-patient variability. The numbers of cells under all the clusters from each individual donor were counted and used as the response variable in the model. The glm function of the stats package was used to fit the cell numbers against cell clusters, donors, plus a cluster-menopausal interaction term. The quasi-Poisson family was used in the glm function.

A quasi-multinomial F-test was performed to test for differences in cluster frequencies across all the clusters between pre- and post-menopausal samples, which yielded a p-value of 0.007. To test for cluster frequency differences for each individual cluster, we compared the cell numbers of that cluster with the aggregated cell numbers of all the other clusters across all the donors. Quasi-binomial generalized linear models were fitted and quasi-binomial F-tests were performed for each cluster separately. The p-values are 0.040 and 0.032 for cluster 1 and cluster 2, respectively, indicating these two clusters have significantly different sizes between pre- and post-menopause conditions after accounting for inter-patient variability. Sizes are not significantly different for other clusters. The R code to reproduce the differential abundance analysis is provided in the files NormEpi.R and NormTotal.R (Table 2).

### Copy number variation analysis

Copy number variation (CNV) analysis was performed using inferCNV of the Trinity CTAT Project (https://github.com/broadinstitute/inferCNV), which compares gene expression intensity across genomic locations in the tumor or lymph-node samples with those in a normal reference sample. The single-cell RNA expression profile of a normal breast total cells sample (N-0372-total) was adopted as a reference for all the CNV analyses presented in the ScBrAtlas study. The results of each CNV analysis were visualized in a heatmap, which showed the relative expression intensities of the tumor samples with respect to the normal reference. For ease of visualization, cells from the same patient within the same cluster were grouped into a single column block, and only the blocks containing more than 100 cells were used in the heatmap. All the column blocks were assigned an equal width in each of the heatmap. The column block annotation of all the CNV heatmaps in this study is available as part of the Figshare deposition, indicating which clusters in which samples were classified as normal or tumor^9^.

## Data Records

Cell Ranger genewise read counts for the 69 scRNA-seq profiles, prior to quality filtering, are available as GEO series GSE161529^7^. Quality filtering thresholds, downstream R data objects storing cell cluster identities and associated R code are available from Figshare9. Specific files available from Figshare are listed in Table 2.

The bulk RNA-seq genewise read counts are available as GEO series GSE161892^24^. The cell-type signature genes generated from the bulk RNA-seq and associated R code are available from Figshare^9^.

## Technical Validation

Technical quality of the 10X single-cell transcriptomic datasets was assessed by examining the number of mapped reads and the number of detected genes (genes with at least one read count mapped to it) for all cells across all the samples (Fig. 2a-b).

**Figure 2.**
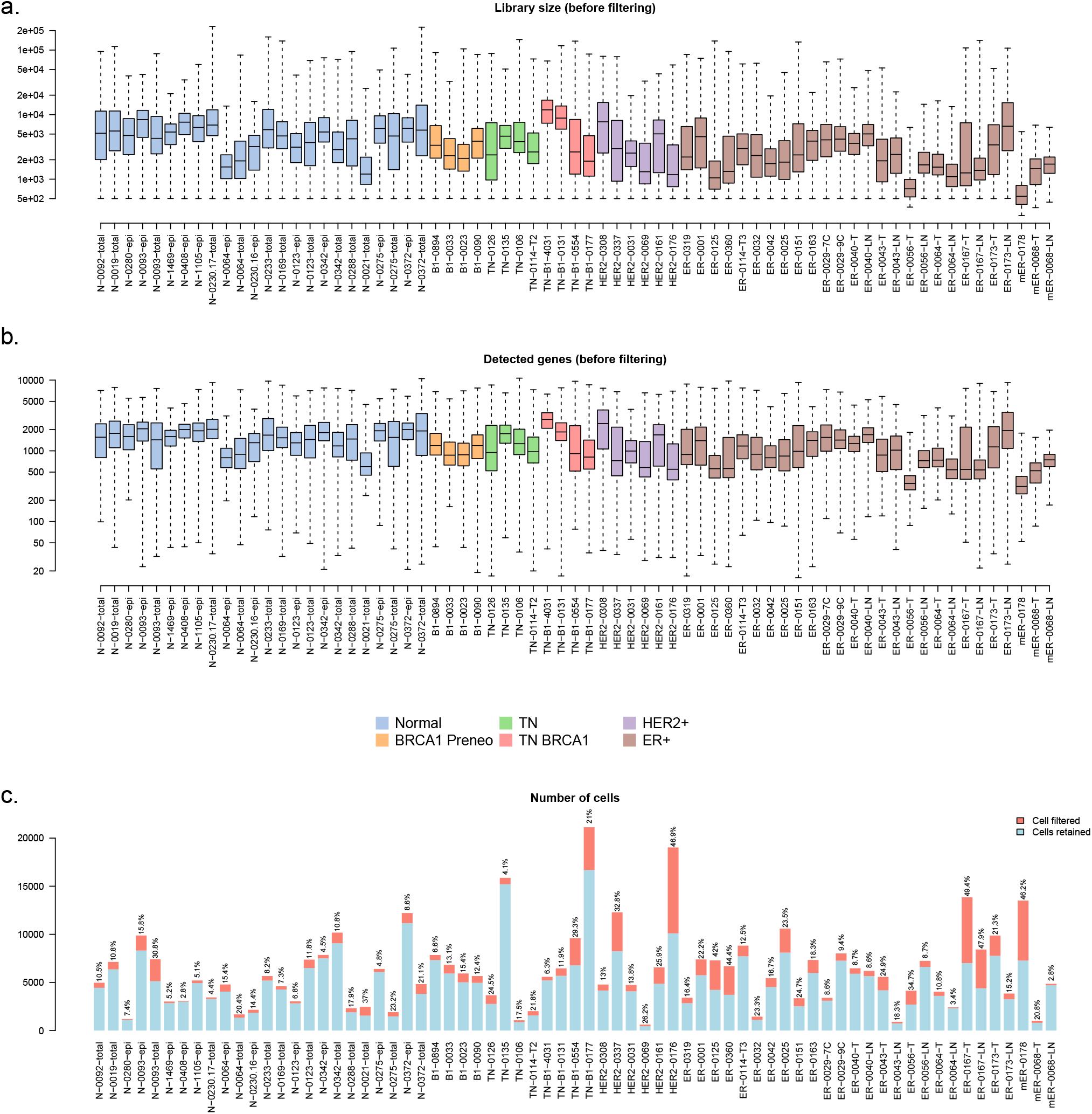
Box plots of (a) the library sizes and (b) the numbers of detected genes for all the cells in each of the 69 samples before filtering. Boxes are coloured by tumor type. (c) Bar plots of the number of cells in each of the 69 samples. The blue segments show the number of cells that are kept after the cell filtering while the red segments show the filtered cells. The proportions of filtered cells are labelled on top of the bars for all 69 samples.

Quality control was performed to remove cells of low quality. Cells with a high proportion of mitochondrial reads or a low number of detected genes were removed. For each sample, an upper limit of library size was also used in combination with an upper limit of number of detected genes to remove potential multiplets. The proportion of cells retained after filtering is 82.2% across all 69 samples, indicating good data quality (Fig. 2c).

Technical quality of the bulk RNA-seq data was assessed using MDS and BCV plots (Figure 3).

**Figure 3.**
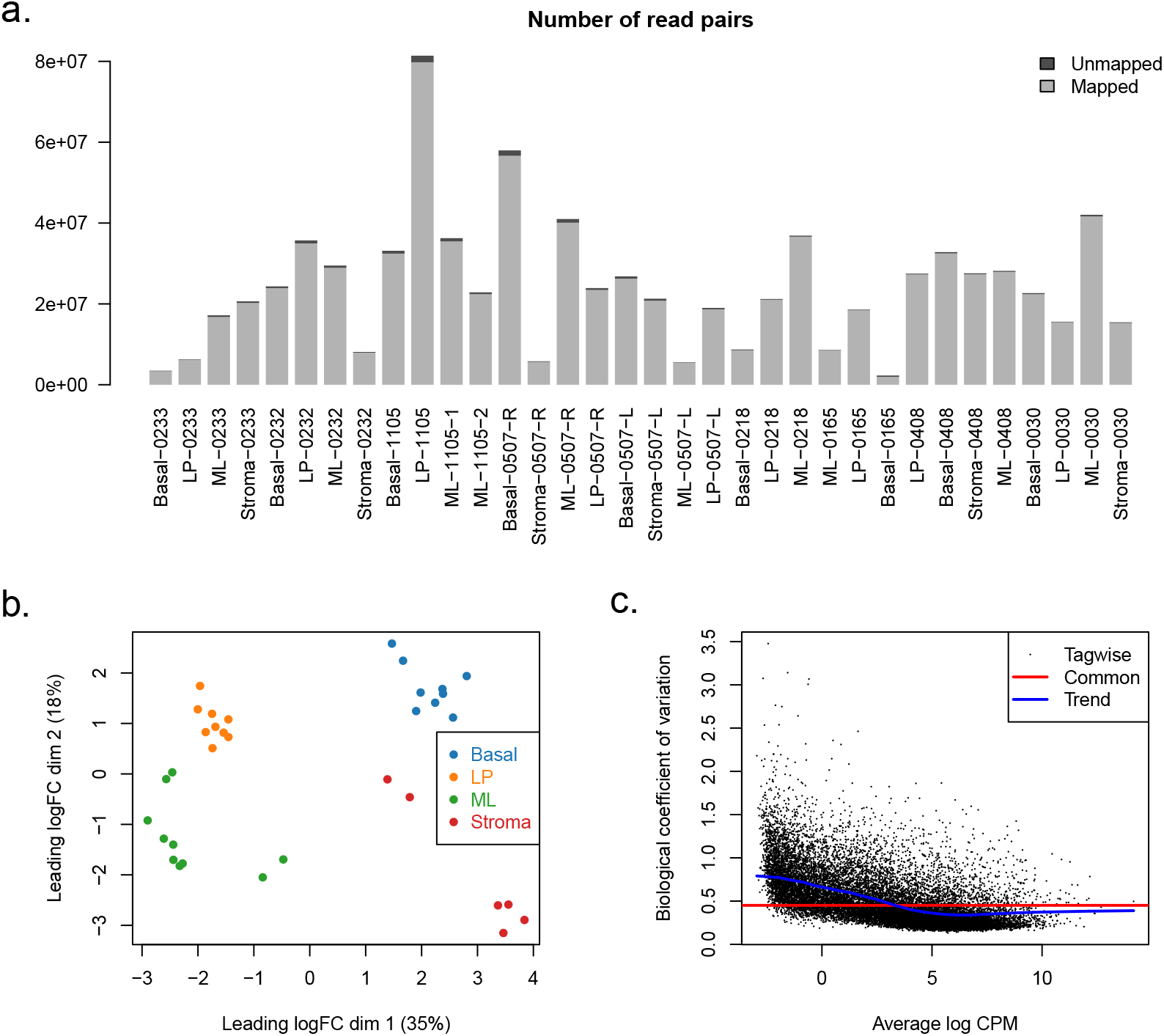
(a) Bar plots of the numbers of read pairs in the human mammary gland bulk RNA-seq samples. The light grey segments represent the mapped read pairs whereas the dark grey segments represent the unmapped ones. (b) MDS plot of all the bulk RNA-seq samples. (c) BCV plot of the bulk RNA-seq data.

## Usage Notes

The code provided may be run using the free R programming environment with Bioconductor and Seurat R software packages https://www.r-project.org. The RDS files may be read using R’s readRDS() function. The Seurat objects allow readers to use and extend the results of the major analyses conducted as part of the ScBrAtlas study. Cell barcodes and Seurat cell clustering information are stored in the meta.data component of each Seurat object.

## Code availability

The R code files provided on Figshare contain complete code and input files for reproducing the analyses of the BrScAtlas study^9^ (Table 2). All the bioinformatics analyses were performed in R 3.6.1 on x86_64-pc-linux-gnu (64-bit) platform, running under CentOS Linux 7. The following software packages were used for the analyses: Seurat v3.1.1, limma v3.40.6, edgeR v3.26.8, pheatmap v1.0.12, ggplot2 v3.2.1, org.Hs.eg.db v3.8.2 and vcd v1.4-5.

## Acknowledgements

This work was supported by the Chan Zuckerberg Initiative (EOSS4 grant number 2021-237445), the National Breast Cancer Foundation (NBCF, IIRS-20-022), Australian National Health and Medical Research Council (NHMRC) grants (#1054618, 1100807,1113133, 1153049); NHMRC IRIISS; the Victorian State Government Operational Infrastructure Support; the Australian Cancer Research Foundation and the Ian Potter Foundation. G.J.L., G.K.S. and J.E.V. were supported by NHMRC Fellowships (G.J.L. #1078730 and 1175960; G.K.S. #1058892; J.E.V. #1037230 and 1102742); Y.C was supported by Medical Research Future Fund (MRFF) Investigator Grant (#1176199).

## Author contributions statement

Y.C. and G.K.S performed bioinformatic analyses, deposited analysis code and data objects, and wrote the article; B.P., J.E.V and G.J.L designed the human breast study and collected data. All authors reviewed the manuscript.

